# ARID5A Is a PERK-Dependent Effector of the Endoplasmic Reticulum Stress Response That Protects Against Steatohepatitis

**DOI:** 10.64898/2026.10.15.759473

**Authors:** Takahito Nishihara, Jasmin Weninger, P. Vineeth Daniel, Hanna L. Erickson, Aditya V. Bhagwate, Amy S. Mauer, Harmeet Malhi

## Abstract

Lipotoxic endoplasmic reticulum (ER) stress is a fundamental feature of metabolic dysfunction-associated steatohepatitis (MASH). ER stress activates adaptive signaling processes that aim to restore ER homeostasis. Yet, depending on the duration, magnitude, and cellular context, sustained ER stress becomes maladaptive and leads to cell death. The regulatory pathways that determine adaptive versus maladaptive ER stress outcomes remain incompletely understood. Here we integrated transcriptomic and chromatin-accessibility analyses in complementary models of proteotoxic and lipotoxic ER stress to identify previously unrecognized regulators of the ER stress response. These analyses identified the dual DNA- and RNA-binding protein AT-rich interaction domain 5A (ARID5A) as an ER stress-responsive gene. ARID5A protein was abundantly expressed in hepatocytes, and ARID5A mRNA and protein were upregulated in palmitate-induced lipotoxic ER stress across multiple hepatocyte models. Using complementary genetic and pharmacological approaches, we demonstrated that ARID5A induction was independent of IRE1α and ATF6α signaling and was attenuated by inhibition of the PERK pathway. To determine the functional relevance of ARID5A in MASH, we employed hepatocyte-directed AAV9-mediated ARID5A overexpression in mice fed a MASH-inducing diet. ARID5A overexpression attenuated steatohepatitis, inflammation, and fibrosis. Together, these findings identify ARID5A as a previously unrecognized PERK-dependent effector of the hepatocyte ER stress response and demonstrate that increasing hepatocyte ARID5A is protective in experimental MASH. This work provides a foundation for defining the mechanisms by which ARID5A regulates hepatocyte stress responses and MASH severity.

## INTRODUCTION

The endoplasmic reticulum (ER) is an essential hub of protein folding, lipid biosynthesis, and calcium homeostasis in hepatocytes. When hepatocytes are exposed to stressors such as lipid overload, a defining feature of metabolic dysfunction-associated steatohepatitis (MASH), the structure and function of the ER are compromised. In response, the unfolded protein response (UPR), a conserved intracellular signaling network, is activated [1, 2]. The UPR is mediated through three primary signaling sensors: protein kinase RNA-like ER kinase (PERK), inositol requiring enzyme 1 alpha (IRE1α), and activating transcription factor 6 alpha (ATF6α) [3]. Each sensor activates downstream programs that collectively restore ER homeostasis in response to cellular stress. While transient activation promotes adaptive restoration of ER function, persistent or unresolved ER stress can be maladaptive, leading to cell death, inflammation, and injury [1]. Thus, the UPR is a dynamic signaling process whose biological output can be determined by the intensity and duration of ER stress.

Each of the three UPR sensors is activated by lipid excess or lipotoxicity in MASH and is linked to diverse metabolic and pathogenic pathways [4–6]. IRE1α activation can regulate steatosis via XBP1, regulatory microRNAs, and nitrosylation [7–10]. ATF6α can be directly activated by sphingolipids [11] and has been implicated in MASH-associated hepatocellular carcinoma [11, 12]. PERK activation leads to eIF2α phosphorylation and selective translation of activating transcription factor 4 (ATF4), which activates adaptive programs but has also been implicated in hepatic steatosis [13, 14]. In sustained ER stress, ATF4 induces C/EBP homologous protein (CHOP), which can cooperate with ATF4 in stress-responsive transcription while also contributing to maladaptive outcomes [15, 16]. Despite substantial mechanistic progress, significant gaps remain in our understanding of the regulatory pathways that modulate the ER stress response in MASH. Our recent report demonstrated a lipotoxicity-influenced enhancer element within the S100A11 gene that is further acetylated in lipotoxicity and in MASH, aggravating disease severity [17]. Informed by this recently identified enhancer element that can be licensed in lipotoxicity, we postulated that additional regulatory pathways activated during hepatocyte ER stress may modulate hepatocyte ER stress outcomes.

Therefore, in this study, we utilized ER stress models and a multi-omic approach to interrogate the regulatory programs activated by ER stress and identify previously unrecognized transcriptional responses. We discovered that ARID5A, a dual DNA- and RNA-binding protein, is transcriptionally upregulated downstream of the PERK pathway. ARID5A is upregulated in MASH; however, further increasing its expression attenuated experimental MASH. These observations provide a foundation for defining the mechanisms underlying the protective effects of ARID5A.

## METHODS

### Mouse studies

All animal procedures were performed following approval from the Mayo Clinic Institutional Animal Care and Use Committee (IACUC) and were conducted in accordance with established Public Health Service guidelines for the humane care and use of laboratory animals. Six-week-old C57BL/6 mice were purchased from The Jackson Laboratory to permit acclimatization. Mice had unrestricted access to food and water and were housed in standard pathogen-free facilities with 12:12-hour day-night circadian cycles.

### Tunicamycin administration

Tunicamycin was administered to chow-fed adult male mice at a dose of 2 mg/kg by intraperitoneal injection. A stock solution (10 mg/mL) of tunicamycin (T7765, Sigma Aldrich) was prepared in DMSO (276855, Sigma-Aldrich). For injection, the tunicamycin stock solution was further diluted in 150 mM D-glucose. Control mice received the corresponding volume of DMSO diluted in 150 mM D-glucose. Eight hours after the injection, mice were euthanized, blood was collected by cardiac puncture, the liver was flushed with 5 mL of cold PBS, excised, and weighed. Immediately, 25 mg of fresh liver was placed in cold PBS with 10% fetal bovine serum (FBS) and processed for assay for transposase-accessible chromatin (ATAC) sequencing. In parallel, 25 mg of each liver was flash frozen in liquid nitrogen, later processed for RNA extraction for sequencing.

### Hepatocyte-specific Arid5a overexpression mouse model

Age-matched adult male mice were placed on choline-deficient, L-amino acid-defined, high-fat diet (CDA-HFD, Research Diets #A06071302) to induce steatohepatitis or chow diet (CD) for controls. Mice received an intravenous injection of 1.0 x 10^11^ viral copies of either adeno-associated serotype 9 virus (AAV9) under the thyroxine binding globulin (TBG) promoter expressing murine *Arid5a* (mArid5a) with the woodchuck hepatitis virus posttranscriptional regulatory element (WPRE; AAV9-TBG-mArid5a-WPRE; Vector Builder) or enhanced green fluorescent protein (EGFP)-expressing control virus. Simultaneously with virus injection, the mice were placed on either the CDA-HFD or CD for 2 weeks. Upon completion of 2 weeks of diet feeding, mice were euthanized following a 6-hour fast.

### RNA extraction and RNA sequencing sample preparation

Total RNA was isolated from 25 mg of flash-frozen liver tissue. Samples were homogenized in 500 μL of TRIzol reagent using an RNase-free hand-held pestle, after which an additional 500 μL of TRIzol was added. Following vortexing, 200 μL of chloroform (MilliporeSigma #CX1058) was added, and the samples were vortexed for 15 s and incubated at room temperature for 3 min. Samples were then centrifuged at 12,000 × g for 15 min at 4°C. The upper aqueous phase (425 μL) was transferred to a gDNA Eliminator spin column, and RNA purification was continued using the RNeasy Plus Mini Kit (Qiagen #74134) according to the manufacturer’s instructions. RNA was eluted in 50 μL of RNase-free water. RNA concentration and purity were assessed using a NanoDrop ND1000 (Thermo Scientific). Samples were subsequently diluted 1:10 in RNase-free water and submitted for further analysis.

### RNA sequencing and ATAC sequencing

RNA sequencing data were processed using the Mayo Analysis Pipeline for RNA-seq (MAPRSeq) [18]. Raw sequencing reads were first assessed for quality using FastQC*. High-quality reads were aligned to the mouse reference genome (GRCm38/mm10) using the STAR aligner [19]. Gene-level read counts were generated using FeatureCounts [20] based on reference gene annotations. Differential gene expression analysis was performed using edgeR, which models count data using a negative binomial framework [21]. Prior to statistical testing, lowly expressed genes were filtered to reduce noise, and normalization factors were calculated to account for differences in sequencing depth and library composition across samples using trimmed mean of M-values (TMM) normalization. Genes were considered significantly differentially expressed if they met the criteria of an absolute log2 fold change greater than 1 (|log2FC| > 1) and an adjusted p-value < 0.05 following Benjamini–Hochberg multiple testing correction [22]. Differential expression results were visualized using volcano plots displaying the relationship between log2 fold change and statistical significance.

Approximately 25 mg of liver tissue from each mouse was subjected to ATAC-seq following the Omni-ATAC protocol [23]. The libraries were purified by AMPure XP magnetic beads with double size selections (0.55X and 1.35X), and the size distribution of library DNA was determined by Fragment Analyzer (Advanced Analytical Technologies, Ames, IA) using a High Sensitivity NGS Fragment Analysis Kit (Cat.# DNF-486). Real-time PCR analysis was performed using SYBR Green universal PCR mixes (Bio-Rad, Hercules, CA) to check the enrichment of accessible regions in the library DNAs. The following quality control primers were used: accessible region #1: Fam86fp-F: 5’-GGCATTAGGGCGGGGATAAA-3’, Fam86fp-R: 5’-TAAGACCTGTCTCCTGGGCT-3’; accessible region #2: Ccdc146-F: 5’- GCTCCGCCTCTTAGAAACCA-3’, Ccdc146-R: 5’-GCGATTCTAACACCTGCCCT-3’; inaccessible region: hATAC-C19 inter-F: 5’-GCTTGTCTTTCCCAAGTTTACTC-3’, hATAC- C19 inter-R: 5’-TAGCTGTCGCACTTCAGAGGA-3’. The libraries were sequenced to 51 base pairs from both ends on an Illumina NovaSeq instrument (S1 kit and flow cell; 125x10^6^ read pairs per library) in the Mayo Clinic Genome Analysis Core. ATAC-seq data were processed using the Mayo HiChIP pipeline [24]. Raw sequencing reads were aligned to the mouse reference genome (GRCm38/mm10) using the BWA aligner [25]. Accessible chromatin regions were identified using MACS2 with a q-value threshold of 0.01 [26]. Differential chromatin accessibility analysis was performed using DiffBind [27], which quantifies read counts across consensus peak regions and identifies regions with significant accessibility changes between conditions. Regions were considered significantly differentially accessible if they met thresholds of |log2FC| > 1 and false discovery rate (FDR) < 0.05. To identify transcriptional regulators associated with changes in chromatin accessibility, motif enrichment analysis was performed on differentially accessible regions using HOMER [28]. Both known motif enrichment and de novo motif discovery were conducted using HOMER’s curated transcription factor motif database, and enrichment scores were calculated by comparing motif frequencies within target regions against matched genomic background sequences. To further characterize transcription factor binding activity, ATAC-seq footprinting analysis was performed using TOBIAS, which corrects for Tn5 transposase insertion bias and identifies transcription factor binding footprints within accessible chromatin regions [29]. Differential footprinting scores were computed to identify transcription factors exhibiting condition-specific occupancy patterns. Integration of RNA-seq and ATAC-seq datasets was performed using in-house Python scripts. Differentially expressed genes were associated with nearby chromatin accessibility changes by identifying ATAC-seq peaks located within 1 kb of gene transcription start sites (TSS). Genes exhibiting concordant directionality between chromatin accessibility changes and gene expression changes were considered candidate regulatory targets. The relationship between chromatin accessibility and gene expression changes was visualized using scatter plots comparing log2 fold-change values derived from ATAC-seq and RNA-seq analyses, and linear regression was used to evaluate global correlations between accessibility changes and transcriptional output. Expression levels of transcription factors corresponding to enriched motifs were extracted from the RNA-seq dataset and visualized using log2-transformed expression heatmaps. All statistical analyses were performed using R and Python [30], and figures were generated in R using the ggplot2 package [31].

### Cell culture and treatment

Immortalized mouse hepatocytes (IMH) and HepG2 were maintained in Dulbecco’s Modified Eagle Medium (DMEM, Gibco) supplemented with 10% fetal bovine serum (FBS) (Thermo Scientific), 1% penicillin-streptomycin and antimycotic (Gibco #15240062) and an additional 1% non-essential amino acids (NEAA, Corning #07725004). AML12 cell line was maintained in DMEM supplemented with 10% FBS, 1% Insulin-Transferrin-Selenium (ITS, Gibco #41400045), dexamethasone 40 ng/mL and 1% penicillin-streptomycin antibiotics. Cells were maintained at 37°C in a humidified atmosphere with 5% CO_2_. Cells were seeded in 6-well plates, grown to approximately 80% confluence, and treated with the following treatment media for 8 hours, unless specified otherwise. For palmitate-mediated lipotoxicity experiments, a stock solution of 80 mM palmitate (Sigma-Aldrich #P0500) was prepared in molecular grade isopropanol (Sigma-Aldrich #I9516) and further diluted to a final concentration of 400 μM in 2.5% FBS-containing media and supplemented with 1% bovine serum albumin (BSA) as previously described by us [32]. For tunicamycin-mediated proteotoxicity experiments, a stock solution was prepared in DMSO and diluted to keep the final concentration of DMSO at 0.1% in the treatment medium comprised of DMEM+2.5%FBS+1%BSA. SV40-T antigen immortalized mouse liver cell lines were a gift from Dr. Randal Kaufman (Sanford Burnham Prebys, San Diego, California, USA), and previously described respectively for IRE1α wild-type (IRE1α-WT) IMH cells, IRE1α knockout (IRE1α-KO) cells, ATF6α-WT cells, ATF6α-KO cells, eIF2α WT (SS) cells and homozygous, phosphorylation-resistant eIF2α cells which carry a mutation of serine 51 to alanine (AA). Inhibitor experiments were performed as follows: cells were incubated with 1.0 μM GSK2606414 (MedChemExpress #HY-18072) for an hour before replacing with treatment media that contained palmitate or tunicamycin along with the respective inhibitors. The volume of vehicle was identical for each inhibitor and included in the vehicle control conditions. Upon completion of treatment, cells were washed with PBS and collected mechanically by scraping in PBS. Cell suspensions were transferred to 1.5 mL microcentrifuge tubes and spun at 2,400 x g for 5 minutes at 4°C to pellet cells which were lysed in TRIzol or RIPA buffer for RNA or protein extraction, respectively.

### Primary mouse hepatocytes

Mice were anesthetized with ketamine and xylazine administered intraperitoneally. The abdomen was opened and vascular structures were identified. The portal vein was cannulated. The liver was perfused with an EGTA solution followed by a collagenase type II solution (Gibco #17101-015); hepatocytes were isolated by gentle agitation and pelleted by centrifugation at 50 x g for 2 minutes. Isolated hepatocytes were resuspended in serum-free Dulbecco’s Modified Eagle Medium and purified using Percoll® [33]. Cells that were 97 to 100% viable were plated on collagen-coated plates in William’s media (Gibco #12551032) supplemented with 10% fetal bovine serum, antimycotic and antibiotic, and ITS at 9.5 x 10^4^ cells/cm^2^. After overnight attachment, the plates were gently washed with PBS and cells were placed in treatment media. After the treatment, cells were harvested for respective analyses.

### RNA extraction and qPCR

Cell pellets were homogenized in 500 μL of TRIzol by pipetting. Fifteen to 20 mg of liver tissue was homogenized in 1 mL of TRIzol with a hand-held homogenizer. Total RNA was extracted from the TRIzol lysate using a commercially available kit (Zymo #R2070) following the manufacturer’s protocol, including an in-column DNase digestion. Isolated RNA quantity and quality were evaluated with a NanoDrop ND1000 (Thermo Scientific). One μg of total RNA was reverse transcribed to prepare cDNA using iScript cDNA preparation kit (Bio-Rad #170-8890). Quantitative PCR (qPCR) reactions were run on QuantStudio 6 Flex Real-Time PCR System using the Power SYBR Green PCR Master Mix (Applied Biosystems #43-676-59). Three or more biological replicates were performed for each qPCR assay. 18S and hypoxanthine-guanine phosphoribosyl transferase (HPRT) were used as reference genes for mouse samples. 18S and actin were used as reference genes for human samples. Relative fold change was calculated using the ΔΔCT method. Primer sequences used in this study are listed in Table 2.

**Table 1:** Ranked HOMER motif enrichment analysis of accessible chromatin associated with the integrated transcriptional reponse

| **RANK** | **Best Motif Match** | **P-value** |
| --- | --- | --- |
| 1 | JunB | 1e-434 |
| 2 | Atf2 | 1e-258 |
| 3 | C/EBPα | 1e-112 |
| 4 | Foxa2 | 1e-78 |
| 5 | Nr2F2 | 1e-68 |
| 6 | NF1 | 1e-47 |
| 7 | Atf-4 | 1e-39 |
| **8** | **Arid5a** | **1e-34** |
| 9 | Tcfap2c | 1e-28 |
| 10 | Hnf4 | 1e-25 |

**Table 2:**
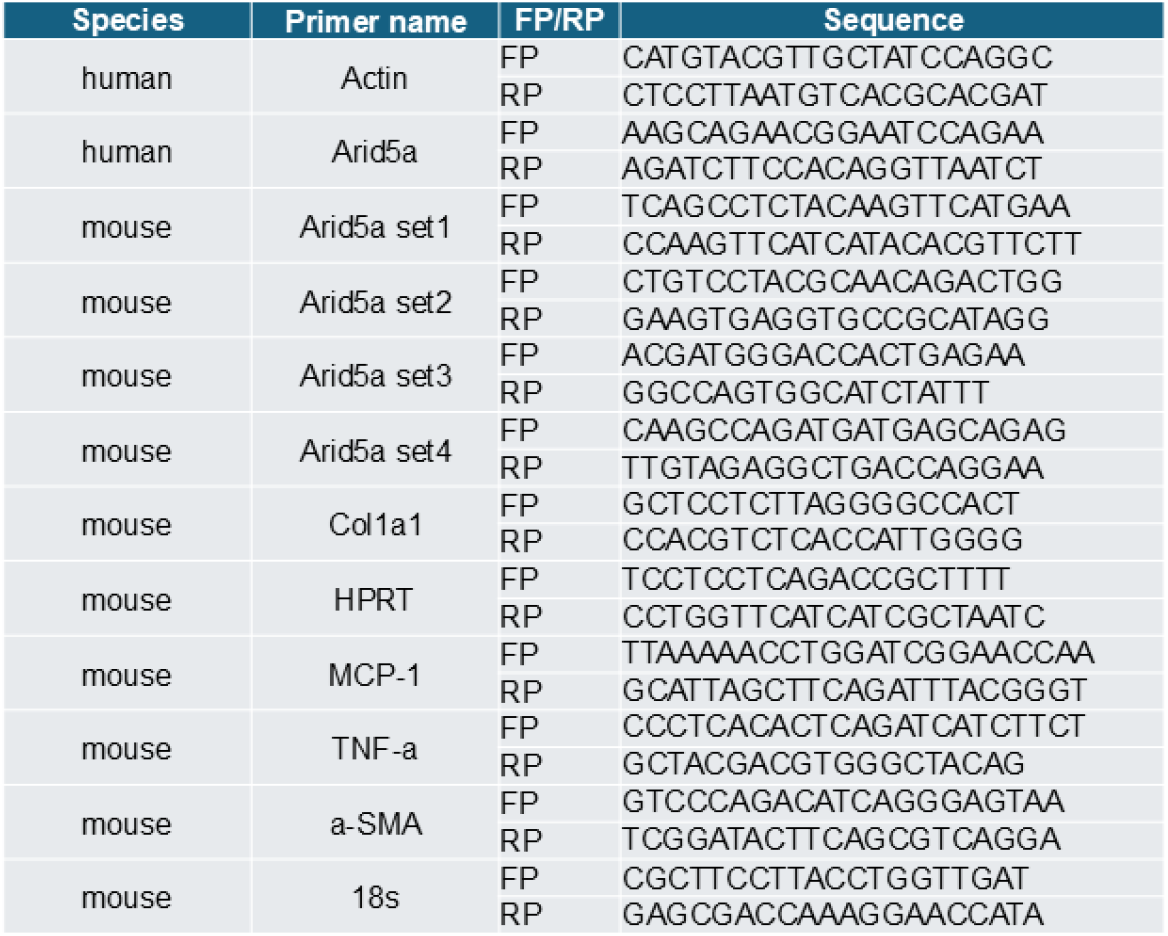
Mouse and human primer sequences

### Western blotting

Whole cell lysates were prepared in RIPA buffer supplemented with protease and phosphatase inhibitor cocktail. Tissue lysates were prepared in T-PER tissue protein extraction reagent (ThermoFisher #78510) supplemented with the same inhibitor cocktail. Total protein was quantified using BCA assay (ThermoFisher #PI23227). Protein from cell (50 μg) or liver (30 μg) lysates was resolved on a 7.5% polyacrylamide SDS gel (Bio-Rad) and transferred onto a 0.22 μm PVDF membrane using Tris-Glycine transfer buffer. Non-specific binding sites were blocked with 5% non-fat dairy milk in TBS-T for an hour at room temperature and incubated with primary antibodies diluted in 5% BSA in TBS-T overnight at 4°C. Next day, membranes were washed, treated with respective horseradish peroxidase-conjugated secondary antibodies diluted to 1:4000 in 5% non-fat dairy milk in TBS-T at room temperature for an hour. For ARID5A, Bullet Blocking One (Nacalai #13779-14) was used as a blocking buffer and Signal Enhancer HIKARI 250 (Nacalai #NU00102) was used as primary and secondary antibody dilution buffer. Following another set of TBS-T washes, signals were visualized using ECL-based chemiluminescence and developed using X-ray films. Western blot densitometry was performed using ImageJ. ARID5A Monoclonal Antibody (Thermo Scientific #MA5-18292) was used for mouse samples, and ARID5A Monoclonal Antibody (Thermo Scientific #MA5-27814) was used for human samples. Alpha-tubulin (Sigma Aldrich #T5168) and vinculin (Sigma Aldrich #V9131) were probed as loading controls.

### Histological analyses

Hematoxylin and eosin (H&E) staining was performed by the Mayo Clinic Histology Core, Arizona. For MAC2 (galectin 3) immunohistochemistry, formalin-fixed paraffin embedded (FFPE) tissue sections were stained with galectin 3 monoclonal antibody (1:400 dilution, ThermoFisher #14-5301-82) using the ABC Immunostaining kit (Vector Laboratories #PK-4001) as previously described by us [34]. Fibrosis was assessed by Picrosirius red staining and quantified by the polarized light microscope method [35].

### *Arid5a* silencing

To knock down *Arid5a*, 400,000 IMH cells were seeded per well of a 6-well plate with Opti-MEM (Gibco #31985070) containing siRNA targeting *Arid5a* (Sigma-Aldrich SASI_Mm02_0034-1523/ARID5A) or a control siRNA (Sigma-Aldrich MISSON siRNA Universal Negative Control) following the Lipofectamine RNAiMAX Reagent (Invitrogen #13778-075) protocol. Four hours after attachment, 10% FBS containing DMEM was added to each well and incubated for an additional 32 hours before collecting cells for isolation of RNA and protein.

### Statistical analysis

All datasets represent mean ± SEM of datapoints obtained from three or more biological replicates. For two experimental groups, two-tailed Student’s t test was used for statistical analyses in GraphPad Prism 10.4.1. Multiple treatment groups were analyzed using ordinary one-way ANOVA, with Bonferroni or Šídák corrections. P < 0.05 was considered statistically significant.

## RESULTS

### Identification of ARID5A as an ER stress-responsive gene

We performed parallel RNA-seq and ATAC-seq in tunicamycin-treated mouse liver tissue, a prototypical model of ER stress [36]. Differential expression analysis of the RNA-seq (Fig. 1A) identified 2,030 genes that were significantly upregulated in tunicamycin-treated livers compared with vehicle-treated livers. Next, within this gene set, to identify robust transcriptional outputs, we integrated ATAC-seq and RNA-seq datasets by intersecting differentially expressed genes with loci exhibiting concordant increases in chromatin accessibility (Fig. 1B). As tunicamycin provides a reductionist platform to selectively resolve ER stress-dependent transcriptional programs, we next investigated these findings in RNA-seq of lipotoxic hepatocytes (Fig. 1C), a model of cellular lipid stress relevant to MASH. Integrated analysis of tunicamycin and palmitate conditions identified a shared transcriptional response across proteotoxic and lipotoxic stress. HOMER motif enrichment analysis of the associated accessible chromatin identified several well-characterized stress-responsive regulators, with ARID5A emerging as the eighth-ranked motif match (p=1x10^-34^; Table 1). ARID5A has not previously been implicated in ER stress, prompting us to investigate its regulation and function in this context. Consistent with our integrated analysis, *Arid5a* was upregulated in the RNA-seq datasets (Fig. 1B, 1C) in parallel with established UPR-responsive genes, including *Ddit3*, *Trib3*, *Atf2*, and *Atf3.* Lastly, we validated these bioinformatic findings by qPCR, demonstrating robust upregulation of *Arid5a* transcripts in tunicamycin-treated mouse livers (Fig. 1D) and palmitate-treated primary mouse hepatocytes (Fig. 1E).

**Figure 1.**
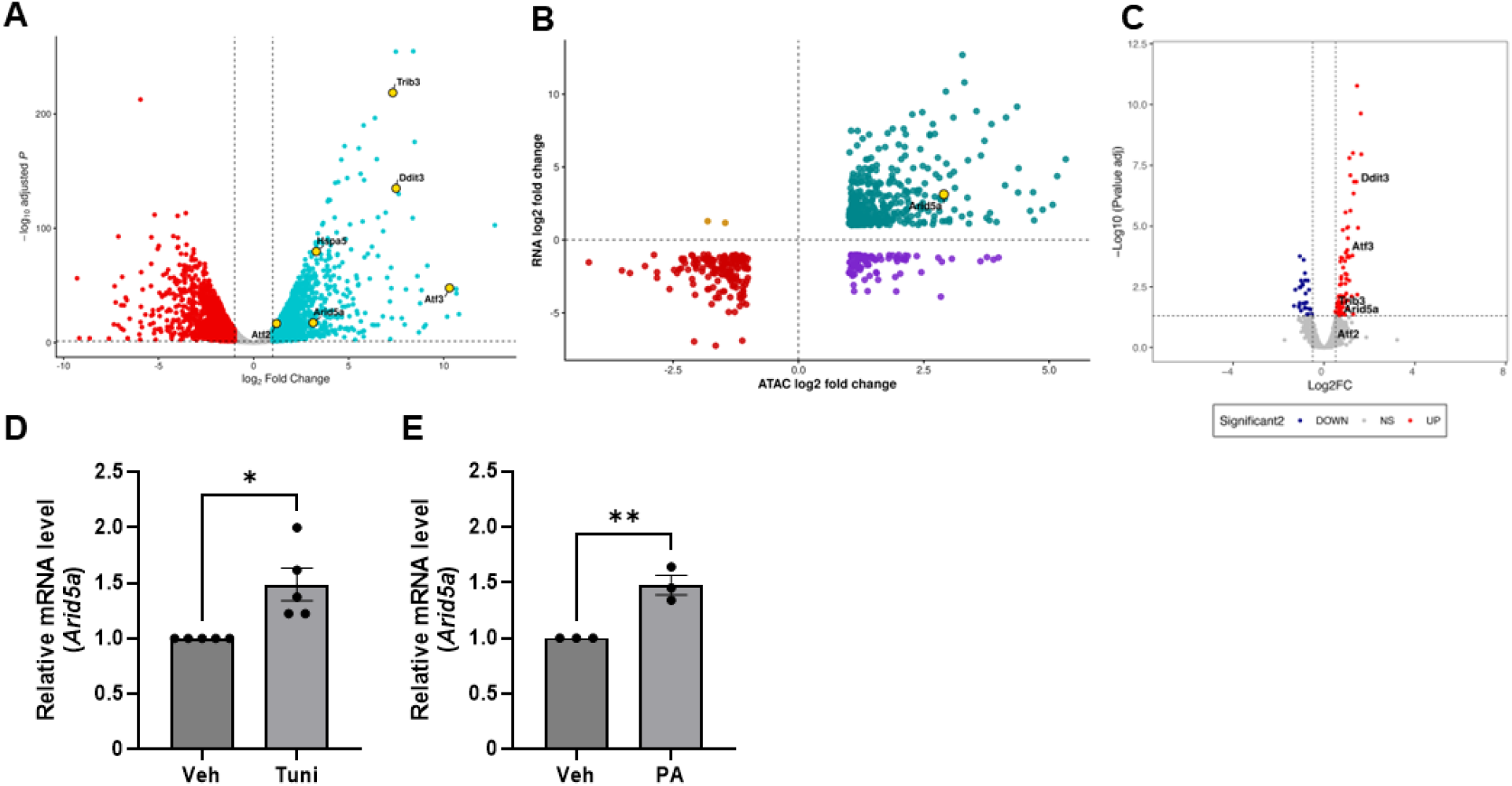
Arid5a is induced by proteotoxic and lipotoxic stress. **(A)** Volcano plot of differentially expressed genes in livers from mice treated with tunicamycin compared with vehicle- treated controls, as determined by RNA sequencing (RNA-seq). The x-axis shows log2 fold change and the y-axis shows −log10 adjusted *P* value. *Arid5a* and selected endoplasmic reticulum (ER) stress-associated genes are highlighted. Genes with |log2 fold change| > 1 and adjusted *P* < 0.05 were considered significantly differentially expressed. **(B)** Four-quadrant plot showing the integration of differential chromatin accessibility determined by ATAC-seq and differential gene expression determined by RNA-seq in tunicamycin-treated mouse livers. The x-axis shows ATAC- seq log2 fold change and the y-axis shows RNA-seq log2 fold change. Genes in the upper-right quadrant (green) show concordant increases in chromatin accessibility and gene expression, whereas genes in the lower-left quadrant (red) show concordant decreases. Genes in the upper-left (yellow) and lower-right (purple) quadrants show discordant changes in chromatin accessibility and gene expression. *Arid5a* is highlighted. **(C)** Volcano plot of differentially expressed genes in primary mouse hepatocytes (PMHs) treated with palmitate compared with vehicle-treated controls determined by RNA-seq. The x-axis shows log2 fold change and the y-axis shows −log10 adjusted *P* value. *Arid5a* and selected stress-responsive genes are labeled. **(D)** *Arid5a* mRNA expression in PMHs treated with tunicamycin (Tuni; 0.5 µg/mL) or vehicle (Veh) for 4 hours, as determined by qPCR and normalized to 18S rRNA (*n* = 5). **(E)** *Arid5a* mRNA expression in PMHs treated with palmitate (PA) (200 µM) or Veh for 8 hours, as determined by qPCR and normalized to 18S rRNA (*n* = 3). Data in (D, E) are presented as mean ± SEM. Statistical significance was determined using an unpaired two-tailed Student’s *t* test. \**P* < 0.05; \*\**P* < 0.01.

#### ARID5A protein is expressed in the liver and in hepatocytes

Given the limited characterization of ARID5A protein expression in the liver, we first validated the specificity of the anti-ARID5A antibody using a knockdown approach, demonstrating corresponding reductions in *Arid5a* mRNA (Fig. 2A) and protein (Fig. 2B, C). Next, we examined ARID5A protein expression in normal human and mouse livers with validated antibodies, which detected ARID5A at the expected molecular weight (Fig. 2D, E). As the liver is multicellular, we employed representative cell lines to determine relative expression in hepatocytes, macrophages, endothelial cells, and cholangiocytes. ARID5A was abundant in hepatocytes (Fig. 2F), though also present at lower levels in the other cell types.

**Figure 2.**
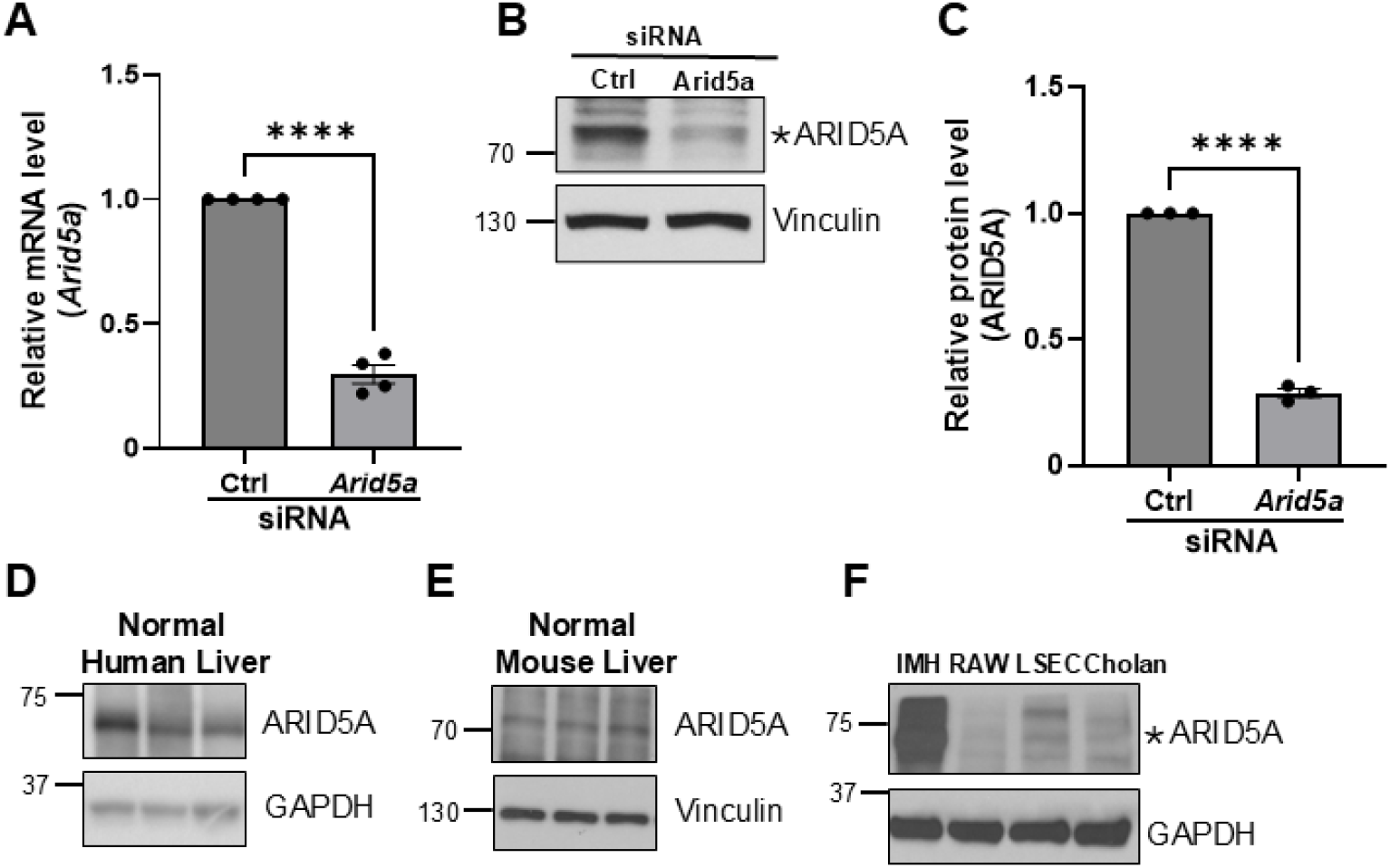
Validation of ARID5A knockdown and characterization of ARID5A protein expression in liver and hepatic cell types. **(A)** *Arid5a* mRNA expression in IMH cells transfected with control (Ctrl) or *Arid5a*-targeting siRNA. *Arid5a* expression was determined by qPCR and normalized to 18S rRNA. **(B)** Representative western blot showing ARID5A protein expression in IMH cells transfected with Ctrl or *Arid5a* siRNA. Vinculin served as a loading control. **(C)** Quantification of ARID5A protein expression shown in **(B)**, normalized to vinculin and expressed relative to controls. Data in **(A, C)** are presented as mean ± SEM (*n* = 4 for **A**; *n* = 3 for **C**). Statistical significance was determined using an unpaired two-tailed Student’s *t* test. \*\*\*\**P* < 0.0001. **(D)** Representative western blot showing ARID5A protein expression in normal human liver samples. GAPDH served as a loading control. **(E)** Representative western blot showing ARID5A protein expression in normal mouse liver samples. Vinculin served as a loading control. **(F)** Representative western blot comparing ARID5A protein expression in immortalized mouse hepatocytes (IMH), RAW 264.7 macrophages (RAW), liver sinusoidal endothelial cells (LSEC), and normal mouse cholangiocytes (Cholan). GAPDH served as a loading control.

### Palmitate treatment upregulates ARID5A expression

We next interrogated ARID5A upregulation in palmitate-induced lipotoxicity in murine and human hepatocytes. In primary mouse hepatocytes, the mouse hepatocyte cell lines AML12 and IMH, and the human hepatocyte cell line HepG2, we found that *Arid5a* transcripts (Fig. 3A-D) and protein levels (Fig. 3E-H) were upregulated following treatment with palmitate. Taken together, these results demonstrate that palmitate-induced lipotoxic ER stress upregulates ARID5A expression in hepatocytes.

**Figure 3.**
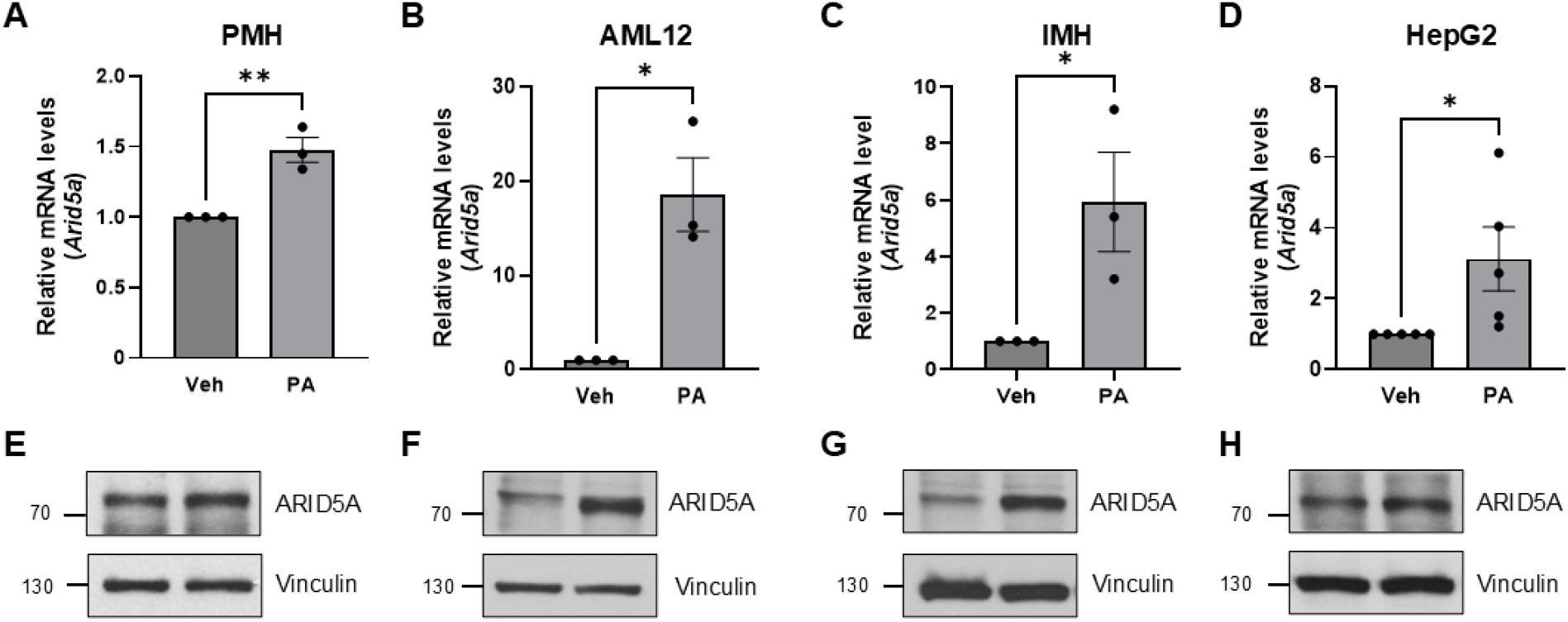
Palmitate treatment increases ARID5A expression across hepatocyte models. (A-D) *Arid5a* mRNA expression in primary mouse hepatocytes (PMHs) **(A)**, AML12 cells **(B)**, immortalized mouse hepatocytes (IMH) **(C)**, and HepG2 cells **(D)** following treatment with palmitate (400 µM) or vehicle (Veh) for 8 hours. mRNA expression was determined by qPCR, normalized to 18S rRNA, and expressed relative to the respective vehicle-treated controls. Data are presented as mean ± SEM; *n* = 3 per group for PMHs, AML12 cells, and IMHs and *n* = 5 per group for HepG2 cells. **(E-H)** Representative western blots showing ARID5A protein expression in PMHs **(E)**, AML12 cells **(F)**, IMH **(G)**, and HepG2 cells **(H)** following treatment with PA or Veh. Vinculin served as a loading control. Statistical significance was determined using an unpaired two-tailed Student’s *t* test. \**P* < 0.05; \*\**P* < 0.01.

### ARID5A is induced downstream of the PERK pathway

ER stress activates the three canonical UPR sensors, PERK, IRE1α, and ATF6α, which activate distinct transcriptional programs [1, 2, 5]. Hence, we interrogated the role of each of the three UPR sensors in regulating the transcriptional upregulation of *Arid5a* in ER stress. We employed a pharmacological inhibitor of the PERK pathway and genetically modified (knockout or loss-of-function) hepatocyte cell lines to identify the UPR sensor necessary for the upregulation of ARID5A in lipotoxic ER stress. IRE1α-knockout (KO), ATF6α-KO, and their respective control cell lines [37, 38] were treated with palmitate. ARID5A protein induction was comparable in the wild-type and KO cell lines (Fig. 4A), suggesting independence from IRE1α and ATF6α signaling. In the presence of the PERK inhibitor, GSK2606414, palmitate-induced upregulation of *Arid5a* was significantly attenuated at both the mRNA (Fig. 4B) and protein (Fig. 4C) levels. Similarly, ARID5A induction by tunicamycin was preserved in IRE1α-KO and ATF6α-KO cells relative to their respective wild-type controls (Fig. 4D).

**Figure 4.**
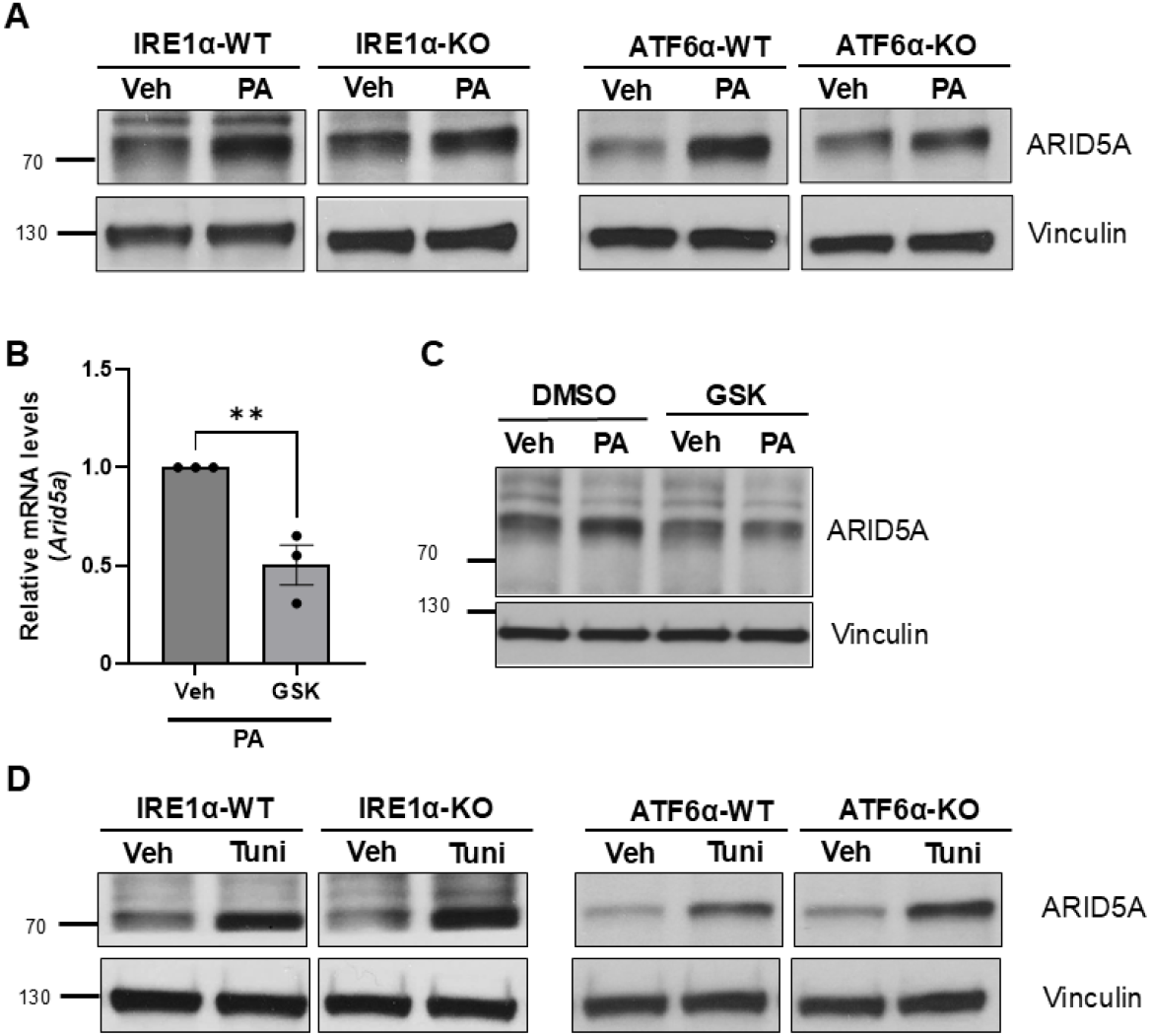
ARID5A induction by lipotoxic and proteotoxic ER stress is independent of IRE1*α* and ATF6*α* and attenuated by PERK inhibition. (A) Representative western blots showing ARID5A protein expression in IRE1α wild-type (IRE1α-WT), IRE1α knockout (IRE1α-KO), ATF6α wild-type (ATF6α-WT), and ATF6α knockout (ATF6α-KO) immortalized hepatocyte cell lines following treatment with palmitate (PA) (400 µM, 8 h) or vehicle (Veh). Vinculin served as a loading control. (B) *Arid5a* mRNA expression in IMH cells treated with PA in the presence of the PERK inhibitor GSK2606414 (GSK) (1.0 μM) or vehicle. *Arid5a* expression was determined by qPCR and normalized to 18S rRNA. Data are presented as mean ± SEM (*n* = 3). **(C)** Representative western blot showing ARID5A protein expression in IMH cells treated with PA or Veh in the presence of GSK2606414 (GSK) or DMSO control. Vinculin served as a loading control. **(D)** Representative western blots showing ARID5A protein expression in IRE1α-WT, IRE1α-KO, ATF6α-WT, and ATF6α-KO immortalized hepatocyte cell lines following treatment with tunicamycin (Tuni; 0.5 µg/mL) or Veh. Vinculin served as a loading control. Statistical significance in **(B)** was determined using an unpaired two-tailed Student’s *t* test. \*\**P* < 0.01.

### ARID5A overexpression attenuates CDA-HFD-induced steatohepatitis

To determine whether ARID5A plays a role in MASH, we used a gain-of-function approach. We employed a hepatotropic AAV9 to overexpress murine ARID5A (denoted as A5OE) in a rapid MASH model induced by feeding the CDA-HFD diet for 2 weeks [39]. Control mice received an EGFP expression control vector (denoted as EGFP). The CDA-HFD feeding was initiated simultaneously with the tail vein injection (Fig. 5A). We chose a two-week timepoint because steatohepatitis with measurable fibrosis is reproducibly induced by the CDA-HFD at this duration of feeding [39]. In terminally collected liver tissue, we noted a significant increase in *Arid5a* expression by qPCR in the A5OE groups regardless of diet (Fig. 5B). Under CD conditions, A5OE resulted in a modest, but non- significant, reduction in serum ALT levels compared with EGFP controls (Fig. 5C). Histological assessment of the cardinal features of MASH in H&E-stained liver sections demonstrated that the CDA-HFD EGFP group developed significant steatohepatitis compared with the CD EGFP group, and ARID5A overexpression (CDA-HFD A5OE) significantly attenuated steatohepatitis (Fig. 5D), fibrosis (Fig. 5D, E), and inflammatory macrophage accumulation (Fig. 5D, F). In keeping with the histological improvement, we observed a reduction in inflammatory (Fig. 5G) and fibrogenic (Fig. 5H) gene expression in CDA-HFD A5OE mouse livers in comparison to the CDA-HFD EGFP group. The CD-fed controls demonstrated no liver injury and normal liver histology in both the EGFP and A5OE groups; thus, there were no apparent adverse effects of ARID5A overexpression.

**Figure 5.**
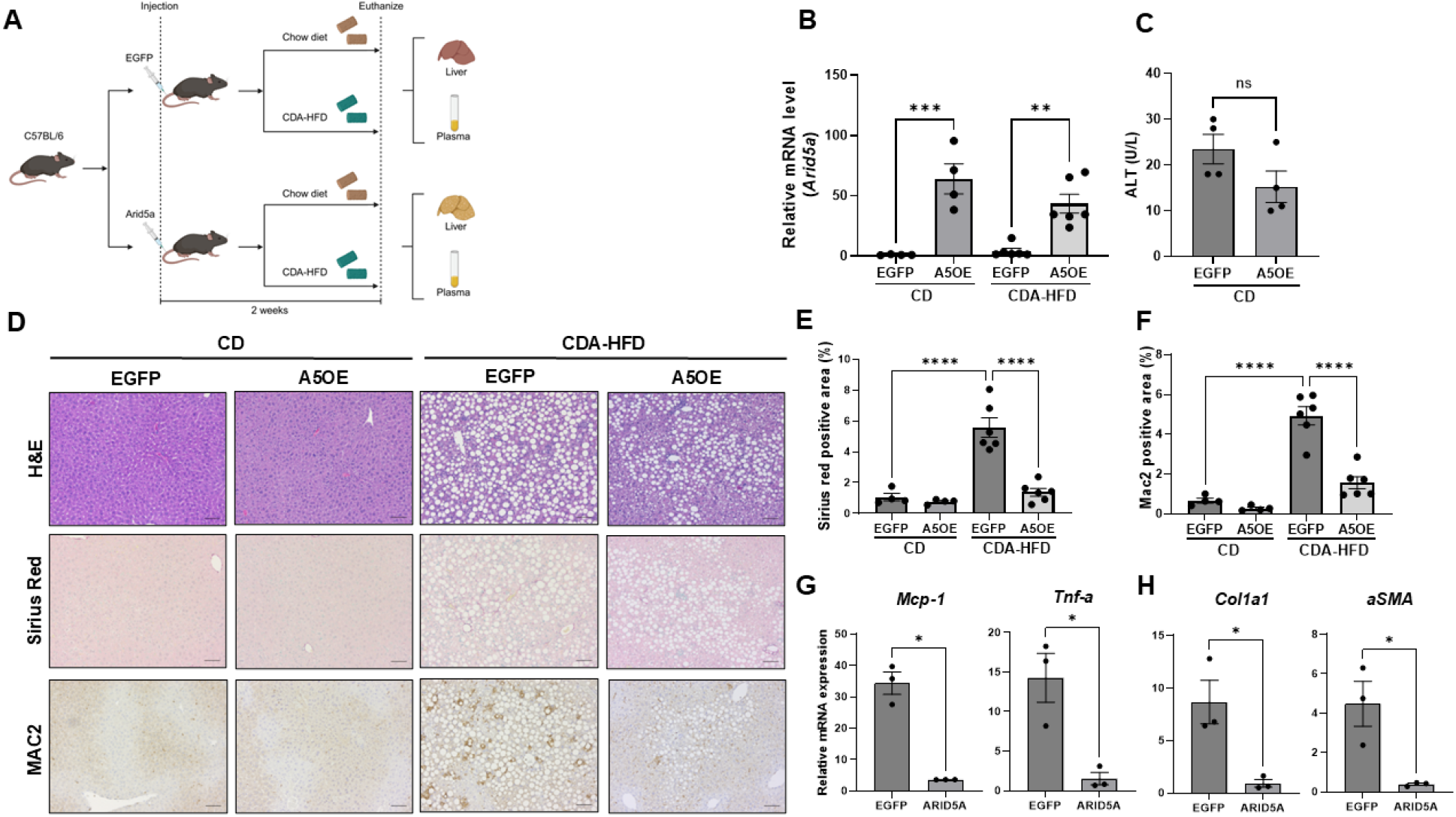
Hepatocyte-specific ARID5A overexpression attenuates CDA-HFD-induced steatohepatitis. **(A)** Schematic design of the hepatocyte-specific ARID5A overexpression study. C57BL/6 mice were injected with control virus expressing enhanced green fluorescent protein (EGFP) or adeno-associated virus (AAV) expressing Arid5a (A5OE) and subsequently fed chow diet (CD) or choline-deficient, amino acid-defined high-fat diet (CDA-HFD) for 2 weeks. Liver and plasma were collected at the experimental endpoint. CD-EGFP (n = 4), CD-A5OE (n = 4), CDA-HFD-EGFP (n = 6), and CDA-HFD-A5OE (n = 6). **(B)** Hepatic *Arid5a* mRNA expression was determined by qPCR and normalized to *Hprt*. **(C)** Plasma alanine aminotransferase (ALT) levels in CD-fed EGFP and A5OE mice. **(D)** Representative liver sections stained with hematoxylin and eosin (H&E), Sirius Red, or MAC2 from EGFP and A5OE mice fed CD or CDA- HFD. Scale bars, 50 µm. **(E)** Quantification of Sirius Red-positive area in liver sections from EGFP and A5OE mice fed CD or CDA-HFD. **(F)** Quantification of Mac2-positive area in liver sections from EGFP and A5OE mice fed CD or CDA-HFD. **(G)** Hepatic mRNA expression of the inflammatory markers *Mcp-1* and *Tnf-α* in CDA-HFD-fed EGFP and A5OE mice, as determined by qPCR. **(H)** Hepatic mRNA expression of the fibrosis markers *Col1a1* and *αSMA* in CDA-HFD- fed EGFP and A5OE mice, as determined by qPCR. Gene expression in **(G, H)** was normalized to *Hprt*. Data are presented as mean ± SEM. Each data point represents one mouse. Statistical significance was determined by two-way ANOVA with Šidák’s multiple-comparison test for comparisons between EGFP and A5OE groups under CD and CDA-HFD conditions, and by unpaired two-tailed Student’s t test for comparisons restricted to two groups. *P < 0.05; **P < 0.01; ***P < 0.001; ****P < 0.0001; ns, not significant.

## DISCUSSION

In the present study we identify ARID5A as a previously unrecognized protective effector of the ER stress response in hepatocytes, downstream of the PERK pathway, and establish its functional relevance in MASH. We report that: i) ARID5A is upregulated in ER stress, ii) this upregulation is PERK-dependent, and iii) in a murine model of MASH, overexpression of ARID5A attenuates liver injury, inflammation, and fibrosis, supporting a protective role for hepatocyte ARID5A in MASH and raising the possibility that augmenting its expression may have therapeutic potential.

We identified ARID5A by integrating chromatin accessibility and transcriptional profiling under proteotoxic ER stress induced by tunicamycin and validating the findings in palmitate- induced lipotoxic stress in hepatocytes. ARID5A induction was conserved across mouse and human hepatocyte models. Both the protein levels and mRNA abundance of ARID5A were increased, suggesting transcriptional upregulation. Previous work has shown that *ARID5A* expression is transcriptionally and post-transcriptionally regulated in the context of inflammatory signaling, notably downstream of NF-κB in immune cells [40]. However, this is the first description of ARID5A regulation in hepatocytes and the first to connect ARID5A to ER stress signaling. We next focused on elucidating the mechanisms leading to the transcriptional upregulation of *Arid5a* transcripts in ER stress and the functional consequences of further increasing its expression.

We investigated the role of each of the three proximal UPR sensors in regulating the expression of *Arid5a.* By employing complementary chemical and genetic approaches, we identified the role of the PERK pathway in the upregulation of *Arid5a*. PERK activation leads to phosphorylation of eIF2α and selective translation of ATF4, a potent transcription factor that in turn induces CHOP transcription. Bioinformatic analysis further identified several ATF4 and CHOP binding sites in the promoter regions of murine and human *Arid5a*, suggesting direct transcriptional regulation by these transcription factors. In future studies we will investigate the individual and combinatorial roles of ATF4 and CHOP in mediating the upregulation of *Arid5a* in ER stress as described for other ER stress-responsive genes [41].

ARID5A is a dual RNA- and DNA-binding protein. Identified as modulator recognition factor 1 (MRF1), with high expression in the aorta, heart, skeletal muscle, and liver, it was isolated as a repressor of estrogen receptor alpha in vascular cells [42]. In contrast, in chondrocytes, ARID5A was reported as an activator of *Col2a1* gene promoter [43]. It was also reported to interact with Sox9 and significantly increase its activity in chondrocytes. A role in adipose tissue homeostasis was identified in global *Arid5a* knockout mice [44]. These mice develop severe obesity in adulthood due to persistent expression of peroxisome proliferator-activated receptor gamma 2 (Ppar-γ2). In contrast, global overexpression of ARID5A protects against high-fat diet- induced obesity [44]. Informed by these observations, we overexpressed ARID5A in a murine dietary MASH model. In CDA-HFD-induced MASH, overexpression of ARID5A attenuated liver injury, inflammation, fibrosis, inflammatory macrophage accumulation, and expression of inflammatory and fibrogenic genes. Altogether, these findings support a protective role for hepatocyte ARID5A in ER stress. Taken together with the observed upregulation in ER stress, we propose a model in which the upregulation of hepatocyte ARID5A may be compensatory, albeit insufficient to mitigate ER stress. Alternatively, supraphysiological expression could engage alternative RNA binding targets or enhance transcriptional activity that is beneficial in MASH. Defining these mechanisms will be the focus of future studies.

### Grant support

This work is supported by NIH grant DK111378 (H.M.), the Mayo Foundation (H.M.), the Optical Microscopy and Clinical Cores of the Mayo Clinic Center for Cell Signaling in Gastroenterology (P30DK084567), and the Histology Core of the Mayo Clinic. J.W. is supported by the Walter-Benjamin-Fellowship of the Deutsche Forschungsgemeinschaft (WE 7859/1-1).

Conflict of interest statement: The authors declare that no conflict of interest exists. Word count: 3948 (Introduction, Methods, Results, Discussion)

## ABBREVIATIONS

A5OE: ARID5A overexpression
AAV: adeno-associated virus
ALT: alanine aminotransferase
ARID5A: AT-rich interaction domain 5A
ATAC-seq: assay for transposase-accessible chromatin with sequencing
ATF6α: activating transcription factor 6 alpha
CD: chow diet
CDA-HFD: choline-deficient, L-amino acid-defined, high-fat diet
DMSO: dimethyl sulfoxide
EGFP: enhanced green fluorescent protein
ER: endoplasmic reticulum
FDR: false discovery rate
H&E: hematoxylin and eosin stain
IRE1α: inositol requiring enzyme 1 alpha
IMH: immortalized mouse hepatocytes
KD: knockdown
KO: knockout
LSEC: liver sinusoidal endothelial cell line
MAPRSeq: Mayo Analysis Pipeline for RNA-seq
MASH: metabolic dysfunction-associated steatohepatitis
OE: overexpression
PA: palmitic acid
PBS: phosphate-buffered saline
PERK: protein kinase RNA-like ER kinase
PMH: primary mouse hepatocytes
Ppar-γ2: peroxisome proliferator activated receptor gamma 2
PVDF: polyvinylidene difluoride
qPCR: quantitative PCR
RIPA: radioimmunoprecipitation assay
RNA-seq: RNA sequencing
Scr: scramble control
SDS: sodium dodecyl sulfate
TBG: thyroxine binding globulin
TBS-T: Tris-buffered saline containing Tween 20 Tuni, tunicamycin
UPR: unfolded protein response
WB: western blotting
WPRE: woodchuck hepatitis posttranscriptional regulatory element
WT: wild-type

